# The natural diversity of the yeast proteome reveals chromosome-wide dosage compensation in aneuploids

**DOI:** 10.1101/2022.04.06.487392

**Authors:** Julia Muenzner, Pauline Trébulle, Federica Agostini, Christoph B. Messner, Martin Steger, Andrea Lehmann, Elodie Caudal, Anna-Sophia Egger, Fatma Amari, Natalie Barthel, Matteo De Chiara, Michael Mülleder, Vadim Demichev, Gianni Liti, Joseph Schacherer, Toni Gossmann, Judith Berman, Markus Ralser

## Abstract

Aneuploidy, an imbalance in chromosome copy numbers, causes genetic disorders, and drives cancer progression, drug tolerance, and antimicrobial resistance. While aneuploidy can confer stress resistance, it is not well understood how cells overcome the fitness burden caused by aberrant chromosomal copy numbers. Studies using both systematically generated ^1–5^ and natural aneuploid yeasts ^6–8^ triggered an intense debate about the role of dosage compensation, concluding that aneuploidy is transmitted to the transcriptome and proteome without significant buffering at the chromosome-wide level, and is, at least in lab strains, associated with significant fitness costs. Conversely, systematic sequencing and phenotyping of large collections of natural isolates revealed that aneuploidy is frequent and has few – if any – fitness costs in nature ^9^. To address these discrepant findings at the proteomic level, we developed a platform that yields highly precise proteomic measurements across large numbers of genetically diverse samples, and applied it to natural isolates collected as part of the 1011 genomes project ^9^. For 613 of the isolates, we were able to match the proteomes to their corresponding transcriptomes and genomes, subsequently quantifying the effect of aneuploidy on gene expression by comparing 95 aneuploid with 518 euploid strains. We find, as in previous studies, that aneuploid gene dosage is not buffered chromosome-wide at the transcriptome level. Importantly, in the proteome, we detect an attenuation of aneuploidy by about 25% below the aneuploid gene dosage in natural yeast isolates. Furthermore, this chromosome-wide dosage compensation is associated with the ubiquitin-proteasome system (UPS), which is expressed at higher levels and has increased activity across natural aneuploid strains. Thus, through systematic exploration of the species-wide diversity of the yeast proteome, we shed light on a long-standing debate about the biology of aneuploids, revealing that aneuploidy tolerance is mediated through chromosome-wide dosage compensation at the proteome level.

## Main

Aneuploidy, a genetic abnormality characterized by the gain or loss of chromosome copies, causes disorders such as Down syndrome, is common in tumor cells, and is an important driver of drug resistance and tolerance, particularly in fungal pathogens ^10–12^. Aneuploidy is also associated with virulence and immune escape in parasites such as *Leishmania donovani* or *Giardia intestinalis* ^13^. At the cellular level, aneuploidy can promote survival of transient environmental stresses, such as heat or high pH, by altering the relative copy numbers of specific genes on the aneuploid chromosome ^14,15^. However, how cells deal with the imbalance of large numbers of genes encoded on the aneuploid chromosome(s) is not well understood.

Several of the seminal studies addressing the consequences of aneuploidy were performed with the budding yeast *Saccharomyces cerevisiae* (*Sc*) because of its amenability to systematic manipulation of chromosomal copy numbers, thereby facilitating comparisons of fitness as well as gene expression levels between aneuploids, and between aneuploids and (otherwise) isogenic parental strains ^1,2,5^. Laboratory-engineered aneuploid yeasts exhibit slow growth rates ^1,2^. Such fitness costs were attributed to the dysregulation of transcripts and proteins encoded in *cis* (on the aneuploid chromosome(s)), and/or dysregulation in *trans* (on other chromosomes) that can cause metabolic, replication, osmotic, and mitotic stress ^13,16,17^. Moreover, in laboratory strains, aneuploidy overloads the ubiquitin proteasome system, causing proteotoxic stress ^18^.

Further, these studies raised a key question around the role of dosage compensation – a potential attenuation of the copy number change at the level of the transcriptome or proteome – in aneuploid cells. Systematic analysis of collections of aneuploid strains revealed that changes in chromosomal copy numbers are largely transmitted to the transcriptome and proteome. For instance, a duplication of any of the 16 *Sc* chromosomes in a haploid background (‘Disomes’) results in a 2-fold increase of the average expression level of both transcripts and proteins encoded on the aneuploid chromosome ^1,4^. A similar conclusion was reached with a collection of strains harboring complex aneuploidies produced from the induced-meiosis of triploid or pentaploid strains ^2,5^. However, these two studies reached different conclusions about the buffering of specific proteins on aneuploid chromosomes: in the disomic collection, components of macromolecular complexes were attenuated ^1,4^, whereas in the induced-meiosis collection no significant protein complex-specific dosage compensation was observed ^2^. As the role of dosage compensation is fundamental to our understanding of the biological basis and consequences of aneuploidy, these divergent results were intensely debated and remain essentially unresolved ^19–21^.

The ease and reduced cost of genome-sequencing facilitated the exploration of the huge biological diversity of *Sc* in the analysis of complex biological processes ^9,22–30^. In the 1011 *Sc* genomes project, budding yeast strains from diverse geographical origins and ecological niches were collected, phenotyped, and genome-sequenced ^9^. 19.1% of these natural isolates carried at least one aneuploid chromosome. This was surprising given the significant fitness costs and often transient nature of aneuploidy in *Sc* lab strains ^1,2,14^. Indeed, only minor growth defects were detected for aneuploid strains from both the 1011 genomes project, as well as in a targeted study of 12 natural aneuploid isolates ^6,7,9^. In order to determine if natural yeasts overcome the fitness costs of aneuploidy through dosage compensation, Hose et al. generated transcriptomes for the 12 natural aneuploids. They found that groups of genes, for example those regulated by feedback control or associated with high-copy toxicity, are attenuated in the natural strains ^6,7^. However, like in laboratory-engineered aneuploid strains, chromosome-wide dosage compensation was not observed ^7,8^.

The proteome of natural strains remained, however, so far uninvestigated. Indeed, while established protocols for genome and mRNA sequencing are applicable on natural strain isolates, their proteomic analysis required us to tackle both biological and technical challenges that we addressed through tailoring cell growth, sample processing, data acquisition and statistical procedures. We then systematically generated proteomes of genetically diverse natural isolates, and processed the strains of the 1011 genomes project ^9^ plus additional strains from a separate natural isolate collection ^31^. Our efforts yielded (to our best knowledge) the first large-scale and precise proteomic dataset of a large natural isolate collection. Further, for a substantial number of the strains (613), we could match these proteomes to the individual strain’s transcriptome and genome. We explored this large multi-omic dataset for a systematic assessment of the pervasiveness of aneuploidy from genome to the mRNA and protein level in natural isolates. Our data confirms that, on the chromosome-wide level, aneuploidy is transmitted unbuffered to the transcriptome ^1–5,8^. However, in sharp contrast to prior studies, we discovered a significant degree of dosage compensation at the chromosome-wide level, and report that proteins that are encoded on the aneuploid chromosomes are systematically buffered. Further, our data associates chromosome-wide dosage compensation to an increased activity of the ubiquitin-proteasome system.

## Results

### A proteomics platform for the precise analysis of large numbers of genetically diverse natural isolates

To adapt high-throughput proteomics for the analysis of natural isolates, we advanced our large-scale proteomic platform originally designed for human plasma ^32–36^. We streamlined the sample preparation workflow to integrate pinning robots for yeast cultivation, as well as liquid-handling stations for sample preparation, enabling the processing of four 96-well plates in parallel (Fig 1a). The mass spectrometric acquisition scheme based on SWATH-MS ^37^ and the non-linear chromatographic gradient (Materials & Methods) were specifically optimized for the precursor m/z distributions and elution profiles of *Sc* tryptic digests to balance throughput and proteomic depth. Moreover, during the project, our Deep Neural Networks-based processing software DIA-NN was continuously developed to reach the processing speed and to include the quality control procedures required to analyze projects in excess of a thousand samples. Indeed, to our knowledge, DIA-NN is, to the best of our knowledge, the only mass spectrometric software specifically addressing proteomic projects of this scale ^33,35^.

**Fig. 1:**
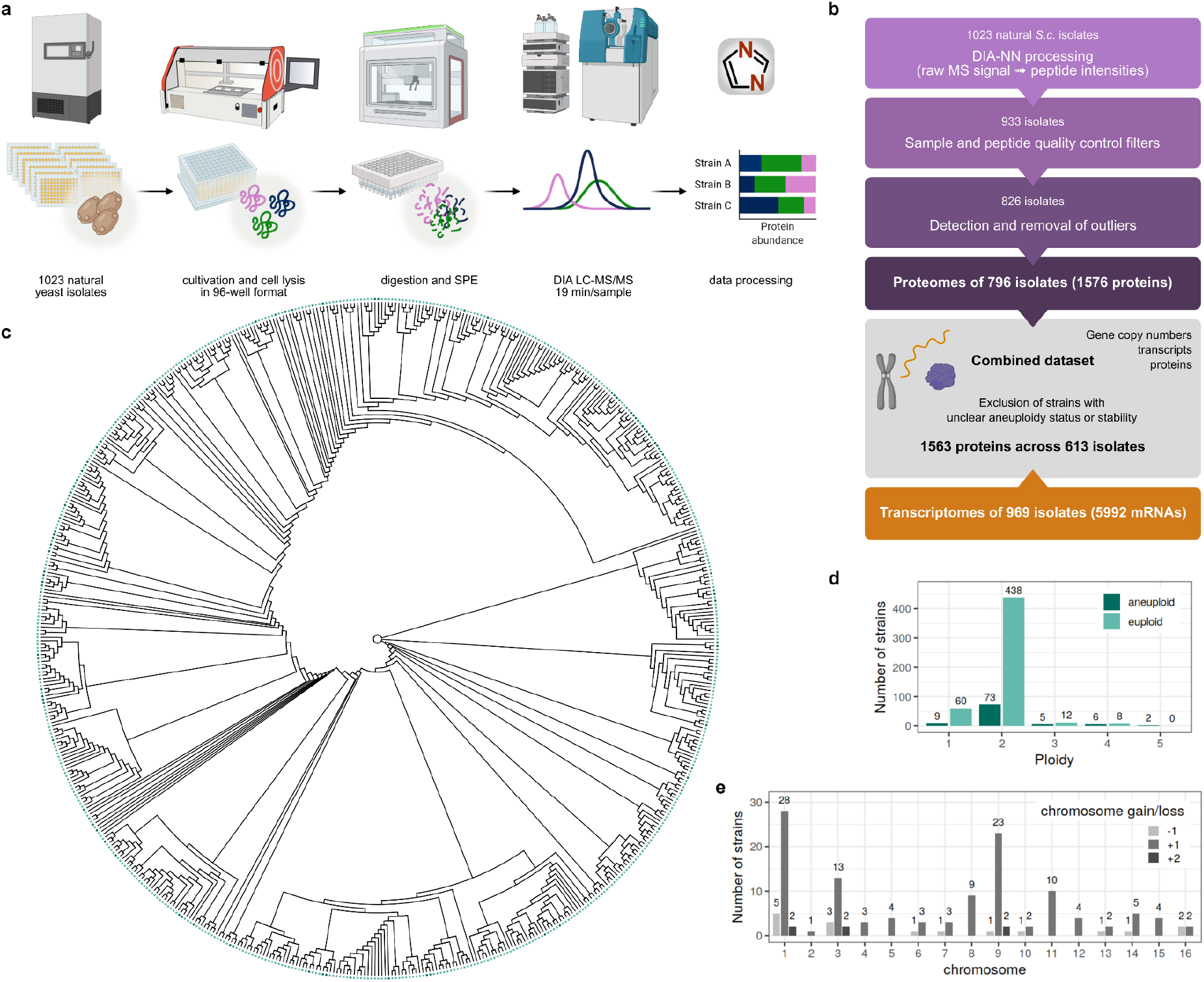
High-throughput yeast proteomics pipeline and assembly of cross-omics dataset for study of aneuploidy in natural yeast strains. (a) *S. cerevisiae* strains in 96-well format were cultivated in synthetic minimal medium. Cells were harvested by centrifugation at mid-log phase and lysed by bead beating under denaturing conditions to prevent non-specific proteolytic digestion. The lysate was then treated with reducing and alkylating reagents and subsequently digested with trypsin. The resulting peptides were desalted by solid phase extraction (SPE) and analyzed by liquid-chromatography tandem mass spectrometry (LC-MS/MS) in data-independent acquisition mode using SWATH-MS. Data were integrated using DIA-NN and subsequently filtered as detailed in Materials and Methods. During cultivation and sample preparation, pinning and liquid handling robots were used to parallelize the workflow. (b) Overview of the proteome processing pipeline and assembly of the combined dataset. Both peptide- and sample-centric quality controls were applied during processing of proteomics raw data. In particular, non-proteotypic precursors, precursors with a high false-discovery rate, as well as samples with low signal and few identifications due to poor growth were removed. (c) Cladogram visualizing the phylogenetic relationship of the 613 isolates of the combined dataset. The tree was pruned from Peter et al. ^9^, keeping the original topology. (d) Numbers of aneuploid and euploid strains per ploidy across the 613 natural isolates included in the combined dataset. (e) Chromosome gains (+1, +2) and losses (−1) across the strains of the combined dataset irrespective of ploidy. For isolates with complex aneuploidies, each aneuploid chromosome was counted separately.

Of the 1023 isolates of the strain library, 933 grew in 96-well microtiter plates and the synthetic minimal medium used to control the metabolic conditions during cultivation ^38^. To control for data quality and to correct for any remaining batch effects, we processed 77 quality control (QC) samples alongside the digests created from the natural strains (Fig S1a, Materials and Methods). Taking advantage of the large number of samples and the low number of missing values, we applied several strict filtering steps to ensure that the dataset from which any conclusions were derived exhibited maximal technical quality, whilst remaining of sufficient size to address the impact of aneuploidies on protein expression across all 16 *Sc* chromosomes and across the genetically diverse strains (Fig 1b). These filters included false discovery rate filters both at the peptide and protein level, filters on precursor numbers, total signal and signal-to-noise ratios, and a filter on precursor variability in quality control samples. Owing to the acquisiton stability of our proteomic technology, we obtained a sufficiently deep proteome even when considering only precursors that were quantified in at least 80% of the 933 isolate proteomes measured, resulting in less than 4% of missing values across all samples (Fig S1b, S1c, Materials and Methods). For the final filtered dataset, we hence retained 1576 protein quantities across 796 strains (SI Table 6). The quantification was precise with a median coefficient of variation (CV) of 14.2% in control samples across all samples processed (Fig S1b). This detected technical variation was much lower than the biological signal (median CV of 31.1%), indicating that we captured proteomic differences across the ∼800 natural strains (Fig S1b).

### Occurrence and frequency of aneuploidy in a natural strain collection

To study the pervasiveness of aneuploidy in natural isolates, we matched the acquired proteomes to annotated genome sequences ^9^ and transcriptomes that were obtained for the natural isolate collection (Materials and Methods). The resulting combined dataset encompassed genomes, transcriptomes, and proteomes for 613 natural strains (Fig 1b), including 518 euploid isolates (haploid, diploid, or polyploid) and 95 aneuploid isolates (Fig 1 c, d). The remaining 183 strains are contained in our proteome dataset, but were not considered for the analysis of aneuploidy because our filtering criteria indicated that there could be a potential mismatch between genome annotation and either aneuploidy annotation^9^ or the mRNA expression profile (Fig 1b, SI Tables 3, 4, Materials and Methods). Such mismatches could emerge if an isolate is not isogenic (i.e. a mixture or two or more strains), but could also emerge due to pipetting errors during the genome sequencing or transcriptomics pipelines, or could be caused by an unstable karyotype. Importantly however, the exclusion of these strains did not bias the dataset for the study of aneuploidy. The relative occurrence of aneuploidies in the filtered and matched dataset was similar to that in the full 1011 genome sequence dataset ^9^.

Of the 518 euploid strains with matched genome, transcriptome and proteome, most were haploid (11.5%) or diploid (84.6%). In the 95 data-matched aneuploids, chromosome gains were much more common (88.4%) than chromosome losses (11.6%). Moreover, aneuploidies were more frequent in strains of higher overall ploidy, with aneuploid strains making up approximately 14% of all haploid and diploid strains, almost 30% of the triploid, and approximately 40% of all tetraploid strains (Fig 1d). Furthermore, the combined dataset covered aneuploidies of all 16 *Sc* chromosomes, with chromosomes I, IX, and III being most frequently aneuploid (25.4%, 18.8% and 13.0%). In contrast, chromosomes II, IV, X, and XIII were amplified only in one to three strains (Fig 1e). Further, 26 strains of the combined dataset contained complex aneuploidies – isolates with multiple chromosome gains, losses or combinations thereof (SI Table 5).

### High-throughput proteomics reveals chromosome-wide dosage compensation in natural strains

First, we assessed how aneuploidy impacted mRNA and protein expression levels by visualizing mRNA and protein levels per chromosome with a heatmap that highlights changes in medians of normalized log_2_ gene expression relative to the change in karyotype (Fig 2a). From the graphical analysis, it is evident that aneuploidy in the genome is transmitted to the transcriptome, as described previously for lab-engineered strains ^1,4,5^, selected wild isolates ^6–8^, and even a collection of aneuploid strains that evolved spontaneously and under minimal selection during mutation accumulation experiments ^3^. Since we studied chromosome- and species-wide mRNA expression distributions instead of performing gene-by-gene comparisons, this result should not be confused, and is compatible, with the situation that the expression of specific individual genes is dosage-compensated at the transcriptome level ^6^. Surprisingly, however, the proteomic response of the natural aneuploids was weaker than the transcriptomic response (generally “dimmer” colors in the proteome versus the transcriptome or genome heatmaps, respectively) (Fig 2a). As chromosome-wide dosage compensation would represent a contrast to previous observations made in lab strains ^1,2^, we asked if this discrepancy could be due to technical differences. Indeed, early-generation proteomic techniques were less precise in quantifying smaller expression changes, especially over large numbers of samples or genes ^20^. We therefore measured the proteomes of the disomic strains described by Torres et al. ^1^ using our proteomics technology (Fig S1d, e, Materials and Methods). Consistent with their earlier report, both mRNA and protein expression changes of disomic strains closely followed the changes in chromosome copy numbers ^1,4^. Thus, the consistent difference in the buffering of aneuploid proteomes between natural strains and lab strains is not due to differences in proteomics technologies.

**Fig. 2:**
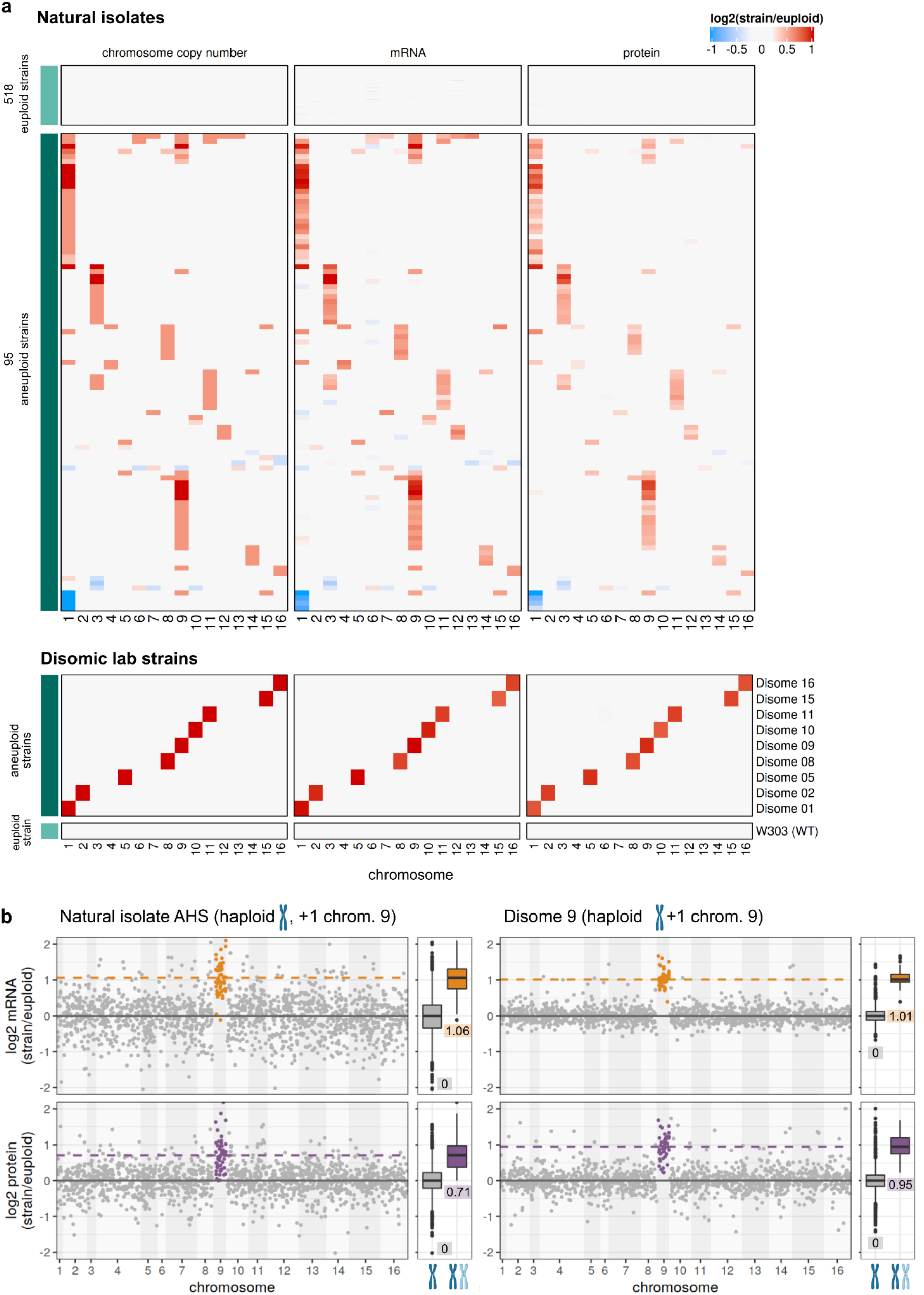
Global transcriptome and proteome profiles across natural and lab-engineered euploid and aneuploid *Sc* strains. (a) Relative chromosome copy numbers, relative median mRNA levels, and relative median protein abundances per chromosome between strains and euploid reference (log_2_ ratios strain/euploid) for natural *Sc* isolates (top panel) and disomic lab-engineered strains (bottom panel). The median across all euploid strains was used as the reference for the natural isolates; the haploid euploid wild-type strain W303 for the disomic strains. (b) Comparison of relative mRNA (upper panels) and protein expression (lower panels) between a haploid natural isolate (AHS) and a haploid disomic lab strain (Disome 9) carrying the same aneuploid karyotype (gain of a single copy of chromosome IX). Genes are sorted by their chromosomal location and their log_2_ mRNA or log_2_ protein ratios are represented as dots. Genes encoded on euploid chromosomes are colored gray, genes located on aneuploid chromosomes are highlighted in orange (mRNA) or purple (protein). The distributions of log_2_ mRNA or protein ratios of all genes encoded on euploid and duplicated chromosomes are shown as boxplots next to the respective chromosome-sorted plots. The median of the distributions is marked with a solid black line within the boxes and displayed below the boxes. Boxplot hinges mark the 25th and 75th percentiles, and whiskers show all values that, at maximum, fall within 1.5 times the interquartile range. For both chromosome-sorted plots and boxplots, relative expression levels are shown between −2 and 2 to improve readability. The solid gray line marks 0 (no change in expression levels), and the dashed colored lines indicate the medians of the aneuploid mRNA or protein expression distributions.

We also compared natural and lab-generated aneuploids with identical karyotypes quantitatively, followed by statistical assessment of dosage compensation across all strains. As a representative example, we compared the natural strain AHS and the lab-engineered Disome 9. Both AHS and Disome 9 are haploid strains with a duplication of chromosome IX (Fig 2b). On the mRNA level, both strains are comparable; the median expression level of a gene encoded on chromosome IX is twice as high in both AHS and Disome 9 compared to the median expression from their respective euploid chromosomes (Fig 2b, upper panels). However, on the proteome level, the strains differ. In Disome 9, the median protein level of genes encoded on the aneuploid chromosome is almost twice as high (log_2_ protein levels = 0.95) as the one of haploid chromosomes. Consistent with previous studies of the disomic strains ^4^, we thus detect no chromosome-wide buffering in the disome. Conversely, in isolate AHS, the median log_2_ protein expression level is only 0.71 (Fig 2b, lower panels), indicating that the increased gene dosage from the aneuploid chromosome is attenuated at the proteome level in the natural strain.

Next, we analyzed the log_2_ mRNA and protein ratios relative to the copy number (CN) change of each chromosome (log_2_ chromosome CN/ploidy) for all natural strains (Fig 3a), and for the disomic strains (Fig 3b). In both the natural strains and the disomes, the distributions of log_2_ mRNA ratios centered around the median predicted by the respective relative chromosome copy number gain, indicating that neither the lab-nor the natural strains exhibited chromosome-wide buffering at the transcriptome level. Similarly, the disome proteomes did not exhibit chromosome-wide buffering, although a subgroup of genes with attenuated protein expression, indicated by a shoulder of the distribution (w = −0.080, p = 1.1 * 10^−3^), was detected, consistent with prior disome analyses ^4^ (Fig 3b). Conversely, in the natural isolates, the log_2_ protein expression distributions of genes encoded on aneuploid chromosomes (log_2_ chromosome CN/ploidy > 0) were shifted down to values closer to that of the euploid state, demonstrating, on average, buffering of proteins encoded by genes on the aneuploid chromosome (Wilcoxon test statistic w between −0.106 and −0.238 with highly significant p-values, Fig 3a). Of note, in cases of chromosome loss, these distribution patterns were mirrored, with transcript levels showing no dosage compensation and protein levels shifting towards the euploid state (Fig S2).

**Fig. 3:**
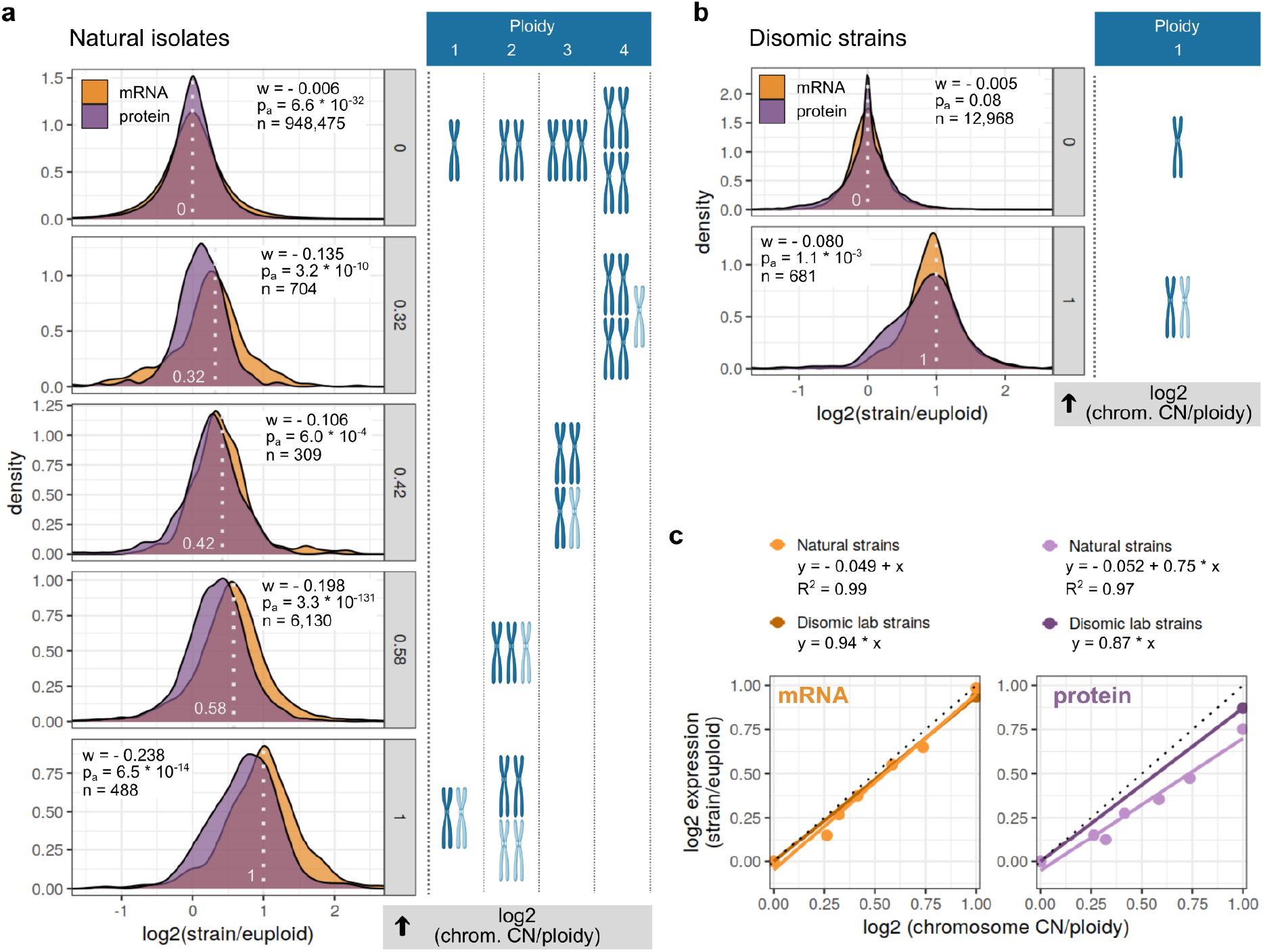
Chromosome-wide dosage compensation of genes encoded on aneuploid chromosomes in natural *Sc* isolates. (a), (b) Distributions of log_2_ mRNA (orange) and protein (purple) ratios across all euploid or aneuploid chromosomes of natural isolates (a) and disomic strains (b). Genes are binned according to the relative copy number (CN) change of the chromosome they are encoded on (log_2_ chrom. CN/ploidy, gray): log_2_ ratios of all genes encoded on euploid chromosomes are summarized in the upper panels (0: chromosome CN equal to ploidy), log_2_ ratios of genes encoded on aneuploid chromosomes are summarized in the lower panels (0.32–1: aneuploid chromosomes in haploid, diploid, triploid, or tetraploid strains). The ploidy of the strains included in the analysis is indicated, and absolute chromosome numbers leading to the shown relative chromosome CN changes when compared to the ploidy of the strain are depicted to the right of each panel, with gained chromosomes highlighted in light blue. The light gray dotted lines and numbers denote the relative chromosome CN change (log_2_ chrom. CN/ploidy) in the density plots. Test statistic (w), adjusted p value (p_a_, Benjamini-Hochberg), and observations (n) for two-sample Wilcoxon tests conducted to compare mRNA and protein distributions for each relative chromosome CN change are shown. For all distributions, relative expression levels are shown between −1.5 and 2.5. (c) Quantification of gene dosage compensation at the mRNA and protein level for natural isolates and disomic lab strains. The median of the log_2_ mRNA and log_2_ protein distributions shown in (a) and (b) is plotted against the relative chromosome CN change (log_2_ chromosome CN/ploidy). The straight lines show the linear regressions for mRNA levels in natural strains (light orange), mRNA levels in disomic lab strains (dark orange), protein levels in natural strains (light purple), and protein levels in disomic lab strains (dark purple). The linear models and, if applicable, R^2^ corresponding to the fitted lines are shown on top of each panel. The dotted black line indicates the expected relative expression levels (log_2_ ratios) in case of no dosage compensation (y = 1*x).

Since the natural yeast dataset included a range of genome sizes from haploid to pentaploid with different chromosome gains and losses (Fig 1d and Fig 1e), we analyzed the relationship between chromosomal copy number and mRNA or protein expression using a relative (log_2_ aneuploid chromosome CN change) range from 0.32 (gain of one chromosome in a tetraploid strain) to 1 (duplication of a chromosome in haploid or diploid strains). On average, the buffering removed about 20%-25% of the aneuploid protein expression level (Fig 3c). In strains with higher ploidies (triploids and tetraploids), chromosome gain results in a smaller relative proteostatic burden and a reduced absolute extent of buffering (Fig 3a, c, lower log_2_ chromosome CN/ploidy numbers). In haploid and diploid strains, gain of one or even two chromosomes leads to larger changes in the overall protein burden the cells experience, and a higher absolute extent of buffering (Fig 3a, c, higher log_2_ chromosome CN/ploidy numbers).

### Chromosome-wide buffering of aneuploidy across diverse natural isolates is attributable to the ubiquitin-proteasome system

To identify mechanistic explanations for the proteome buffering in natural isolates, we took three different approaches. First, we investigated the global proteomic response to aneuploidy in an unbiased manner by performing gene set enrichment analysis (GSEA) of proteins expressed in *trans* in natural aneuploids versus euploids; the KEGG gene set ‘proteasome’ was the most enriched term (Fig 4a). The proteasome was further the only enriched term that was not either related to growth rate (i.e. oxidative phosphorylation, starch and sucrose metabolism), which is also slightly altered in the natural aneuploids (see Peter et al. ^9^, Fig S3a), or related to transcription, which is altered in aneuploids due to the aberrant chromosome number (i.e. nucleotide metabolism, mRNA transport, and transcription, Fig 4a). This result suggested that the aneuploid isolates had higher expression levels of proteins involved in protein degradation, especially structural components of the proteasome.

**Fig. 4:**
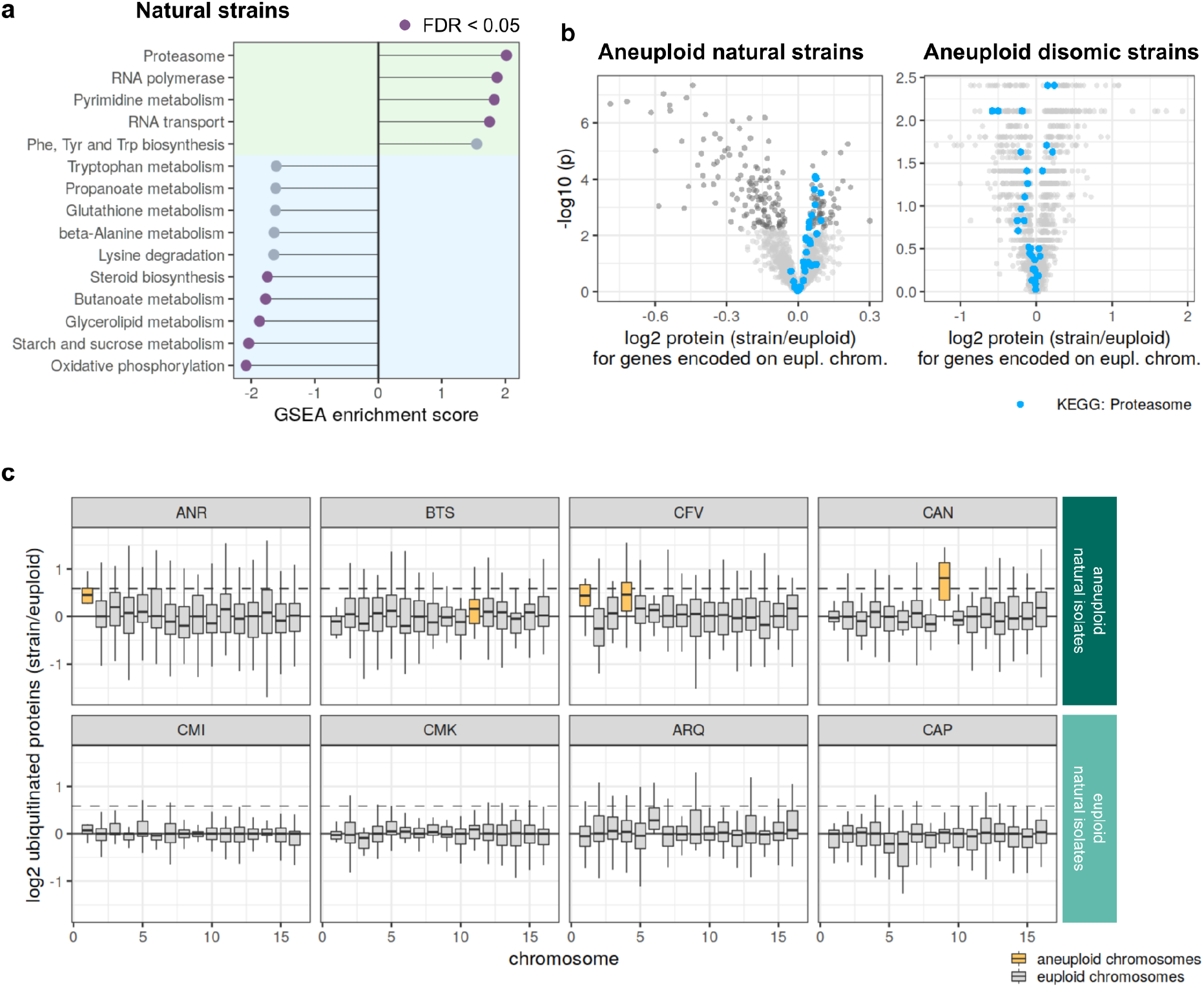
Upregulation and increased activity of the proteasome in natural aneuploid isolates. (a) Gene set enrichment analysis of median log_2_ protein expression ratios (strain/euploid) of genes encoded on euploid chromosomes across all aneuploid natural strains (genes in *trans* of aneuploid chromosomes). Statistically significant enrichment scores (false discovery rate < 0.05) are colored in purple. (b) Volcano plots for natural and disomic strains showing results of non-parametric one-sample Wilcoxon tests comparing the median log_2_ protein ratios to the theoretical gene-by-gene median protein log_2_ ratio across all euploid strains of the respective collection (µ = 0). Proteins with statistically significant differential expression after multiple hypothesis correction (Benjamini-Hochberg) are colored in dark gray. Structural components of the proteasome are highlighted in blue. (c) Chromosome-wide distributions of relative abundances of ubiquitinated proteins in aneuploid and euploid natural isolates. Proteins carrying ubiquitin side chains were used to determine normalized log_2_ protein abundance ratios (comparing expression of each gene per strain with the median expression of the respective gene across the euploid isolates). Chromosomes annotated as aneuploid according to Peter et al. ^9^ are highlighted in yellow, euploid chromosomes are colored gray. The boxplot hinges mark the 25th and 75th percentiles, and whiskers show all values that, at maximum, fall within 1.5 times the interquartile range. Outliers are not shown to improve readability of the distributions.

Second, and equally important, when we compared the proteome of the natural isolates to that of the lab strains, we observed that the enrichment of the UPS genes was specific to the natural aneuploids. An enrichment of the UPS term was neither observed in disomes ^1^ nor in induced meiosis strains ^2^ (Fig 4b, Fig S3b, Fig S4). For example, among the upregulated proteins in the disomes, only five were in the UPS KEGG gene set; among downregulated proteins, 12 were in the UPS KEGG gene set (Fig 4b). This result is consistent with a role for the UPS in buffering the proteomes of natural aneuploid isolates, which does not occur in the lab isolates.

Third, we wanted to directly measure UPS activity in natural aneuploids to determine if it differs between aneuploids and euploids. Because proteasomal inhibitors such as MG132 are not effective in natural yeast isolates due to effluxing ^39^, we applied a recently developed, high-end ubiquitinomics technology based on quantifying the proteolytic remnant of ubiquitinated lysine residues (K-GG remnant peptide profiling) ^40^ and adapted it to yeast samples. The K-GG peptide profiles were recorded for four natural aneuploid isolates selected from the pool of aneuploid strains that had increased proteasome-component protein expression, and each of these strains was matched to a genetically close natural euploid strain as a control (SI Table 5). This revealed stoichiometric enrichment of K-GG-modified proteins encoded on aneuploid chromosomes relative to those encoded on euploid chromosomes (Fig 4c). In parallel, for the same set of four aneuploid and euploid matched isolates, we directly measured proteasome activity in cell extracts. Interestingly, the natural aneuploid isolates degraded Suc-Leu-Leu-Val-Tyr-AMC, a succinyl-coupled (Suc-) model proteasome substrate that releases fluorescent 7-amino-4-methylcoumarin (AMC) when degraded by the 20S proteasome, faster than the matched euploid controls (Fig S3c).

## Discussion

Natural isolate collections have been recognized as powerful tools to study biology beyond selected laboratory-adapted genetic backgrounds ^9,22,23,28,41^, thus supporting the fine mapping of genetic traits and enabling the dissection of the genetic basis of phenotypic variability ^25,29,30,42^. However, thus far, such studies have not been extended to the proteome level, largely due to the technical limitations of proteomic platforms. Here, we present an advance in proteome technology that simplifies the high-throughput acquisition of proteomes from genetically diverse isolates. We applied this technology to a large collection of natural *Sc* isolates, generating highly complete proteomes for 796 strains, thus recording their proteome-wide diversity and determining that chromosome-wide dosage compensation of aneuploidy in natural isolates occurs at the proteome level.

Natural strain collections facilitate a powerful new approach to address complex biological questions. Of the many problems that can be interrogated with our large systematic resource (that will be made publicly available as part of this publication), we herein applied it to investigate the mechanism of aneuploidy tolerance in natural yeast strains ^19,20^. It has long been debated whether dosage compensation contributes to fitness recovery of aneuploids, and here we extend the analysis to a large number of natural aneuploid isolates and to the proteome level ^20^. For 613 out of the 796 strains for which we present proteomic data, the proteomes could be compared to matched transcriptomes and genomes, thus generating, to our knowledge, one of the largest diversity-capturing multi-omic datasets of the current literature. The natural strain dataset complements previous studies using laboratory-generated aneuploid strains by providing general answers to questions that are too complex to be fully answered by relying on individual laboratory strains. Indeed, we discovered a high degree of chromosome-wide buffering of protein levels of genes encoded on aneuploid chromosomes – a feature that had not been observed in aneuploid laboratory strains ^1–5^, or selected wild isolates that had been primarily investigated at the transcriptome level ^6–8^.

Aneuploidy is recognized as a general mechanism that, at least transiently, can render cells stress-tolerant. For example, experimental evolution of *Sc* lab strains in heat or pH stress conditions selected for progeny with chromosome III trisomy or chromosome V disomy, respectively ^14^. Similar examples of “salvage by aneuploidy’’ in *Sc* include rescue of a myosin gene deficiency ^43^, or tunicamycin-induced ER stress that can be countered by chromosome II aneuploidy ^44^. In pathogenic fungi such as *Candida albicans* or *Cryptococcus neoformans*, specific aneuploidies can increase antifungal resistance or tolerance and can drive cross-adaptation to different antifungal drug classes or to antifungals and chemotherapeutics ^45–48^. The increased survival of aneuploid cells under stress conditions is often dependent on one or more key stress response genes that are encoded on the aneuploid chromosome, thus acting through an increased gene dosage, although the slower growth rates of some aneuploids and other general aneuploidy-associated features might confer benefits as well^14.^

Laboratory studies have emphasized the fitness cost incurred by aneuploids when grown in the absence of obvious stresses, although the price of aneuploidy differs depending on the aneuploid chromosomes ^47^; the fitness cost is also more evident in the disomic collection, which must be maintained under G418 selection ^1^, than in the induced-meiosis strains, which were selected for aneuploid stability ^2^. In aneuploids, extra DNA needs to be synthesized, more mRNAs are transcribed and subsequently translated ^49^. As a consequence, altered assembly dynamics of protein complexes with components encoded on the aneuploid chromosome are likely to cause proteotoxic stress ^50,51^ and slow cell growth. Yet the 1011 *Sc* genomes and similar natural yeast strain sequencing projects demonstrated that aneuploidies are common in nature ^9,22^, at least some natural aneuploidies appear to be evolutionarily ancient, and the fitness of natural aneuploids with an extra chromosome is only marginally reduced relative to natural euploid strains ^52^. Thus, it appears that natural strains have found strategies to compensate, to a large degree, for the cost of aneuploidy.

Compensation for the burden of aneuploidy could occur by multiple molecular mechanisms. Our results show, in agreement with previous studies ^1–7^, that mRNA levels are not buffered chromosome-wide, and thus any dosage-compensation must occur post-transcriptionally. Consistent with this hypothesis of post-transcriptional mediation of the burden of aneuploidy, a recent study that analyzed ribosomal footprints in the induced-meiosis aneuploid collection also found no evidence for co-translational buffering of aneuploidy ^5^.

The data we present here for natural isolates point to protein degradation as the major mechanism of dosage compensation at the chromosome-wide level, via the ubiquitin-proteasome system (UPS). This conclusion is based upon several different system-scale approaches. The structural components of the proteasome formed the most enriched gene set of upregulated *trans* genes in natural aneuploids, yet were not induced in the engineered disomes and induced-meiosis aneuploids. Furthermore, the UPS was consistently more active in aneuploids than in euploids, as revealed by deep ubiquitinome profiling and proteasome activity assays with a model substrate. While these system-scale experiments clearly associate chromosome-wide dosage compensation to the UPS, detailed biochemical studies, which are difficult to conduct in large sample series, will be required to clarify the precise molecular interactions that drive UPS activation. Based on these data, we posit that the higher abundance and higher ubiquitination levels of the proteins encoded on the aneuploid chromosome likely lead to their degradation, and that this increased degree of protein degradation requires more UPS activity.

Why were chromosome-wide dosage compensation and upregulation of the UPS not evident in previous studies of aneuploid laboratory strains? Since we recapitulated the observation of little buffering in the disome collection ^4^, the difference cannot be a mere technical issue. Two features of the disome collection likely contribute to the difference in buffering from natural aneuploid isolates. First, the disome collection was constructed in the W303 genetic background, which carries a non-functional *SSD1* allele that affects aneuploid stability ^21^. Second, in the disome collection, the aneuploid chromosomes are lost if they are not maintained under antibiotic selection ^1^. Indeed, when we measured the disomic strains using our proteomics platform, two strains, Disome 12 and Disome 14, had lost their aneuploid chromosome despite antibiotic selection, as evident from their euploid-like protein expression profile (Fig S5). In contrast, the induced-meiosis strains were screened for relative stability of the aneuploid chromosome(s) ^2,5^ and may have adapted partially to their aneuploid state. While we did not detect induction of the proteasome gene set in the five induced-meiosis strains that we analyzed (Fig S4), possibly due to the low number of strains for which proteomic data were previously acquired ^2^, ribosomal profiling suggested that the translation of UPS components was enhanced in these aneuploids ^5^. Proteomic analysis of a larger collection of stable lab-engineered induced-meiosis strains could potentially solve the question whether, and if, to what extent, these strains exhibit dosage compensation.

Based on the data presented here, we propose the following model that explains the differences between lab-generated and natural strains, as a process in which cellular adaptation leads to chromosome-wide dosage compensation. If an aneuploidy provides a selective advantage in specific environmental or stress conditions ^2,12,14,44^, and if this selective advantage outweighs the fitness costs created by the aneuploidy, the aneuploid strains will be selected. These ‘naive aneuploids’ lack chromosome-wide dosage compensation, incur fitness costs, and therefore are transient, as long as the environmental or stress condition is temporary ^1,14^. If environmental conditions continue, selective pressure will favor more stable aneuploid chromosomes; according to our data, this adaptation entails chromosome-wide dosage compensation and increased UPS activity. Indeed, even disomic strains, when evolved *in vitro* under selective antibiotic pressure to maintain their aneuploidy, can accumulate adaptive mutations that affect the UPS and increase protein degradation levels ^18^. Buffering of aneuploidy by dosage compensation has been detected in other species as well. In *Leishmania donovani*, six highly aneuploid strains exhibited no dosage compensation of the transcriptome, while the overall protein level of genes on the aneuploid chromosome was buffered ^53^. In several human cell lines, there is some evidence for chromosome-wide dosage compensation, although the data sets are of moderate size, and the expression differences were too narrow to reach firm conclusions ^20^. Thus, the extent of proteome buffering in organisms that tolerate aneuploidy could be conserved across species.

In summary, our study demonstrates the generation of precise proteomes for a large collection of natural yeast strains, and the matching of these proteomes to the transcriptome and genome of euploid and aneuploid strains. We used the proteomic dataset to explore the natural diversity of yeast in relation to chromosomal aneuploidies; clearly this resource will be useful for addressing many different research questions. We reveal chromosome-wide dosage compensation as a common mechanism mediating aneuploidy tolerance. Furthermore, our results complement observations about the genetic differences between disomic and stable lab-engineered aneuploidies ^21^, shedding light on problems that were heavily debated, and difficult to solve in lab strains and with small sample numbers. Our work highlights the importance of investigating biology beyond selected laboratory models ^41^, and showcases the power of exploring the natural diversity of the proteome to unravel fundamental aspects of cell biology.

## Materials and methods

### Reagents

Unless otherwise noted, reagents were purchased as follows: Bacto Yeast Extract (Cat#212750), Bacto Peptone (Cat#211677), Bacto Dehydrated Agar (Cat#214010), Water (LC–MS grade, Optima, Cat#10509404), acetonitrile (LC–MS grade, Optima, Cat#10001334), methanol (LC–MS grade, Optima, Cat#A456-212) and formic acid (LC–MS grade, Thermo Scientific Pierce, Cat#13454279) were purchased from Fisher Chemicals. Trypsin (Sequence grade, Cat#V511X) was purchased from Promega. D-Glucose (Cat#G7021), Yeast Nitrogen Base (Cat#Y0626), glycerol (Cat#G2025), DL-dithiothreitol (BioUltra, Cat#43815), iodoacetamide (BioUltra, Cat#I1149) ammonium bicarbonate (eluent additive for LC–MS, Cat#40867), yeast nitrogen base without amino acids (Cat#Y0626) and glass beads (acid washed, 425–600 µm, Sigma, Cat#G8772) were purchased from Sigma-Aldrich. Urea (puriss. P.a., reag. Ph. Eur., Cat#33247H) and acetic acid (eluent additive for LC–MS, Cat#49199) were purchased from Honeywell Research Chemicals. 96-well solid-phase extraction plates (MACROSpin C18, 50–450 μL, Cat# SNS SS18VL) were purchased from the Nest Group.

### Yeast strains

The *S. cerevisiae* isolates included in this study were kindly provided by J. Schacherer (Université de Strasbourg, France) and G. Liti (Université Côte d’Azur, France). The library counts 1023 strains in total, of which 997 strains were previously described by Peter et al. ^9^ to be representative of the entire *S. cerevisiae* species. Further 26 strains were originally described by Marsit and Dequin ^54^ and Legras et al. ^31^. The isolates were arranged in a 96-well plate format according to estimated growth rates from solid culture. Aneuploidies in strains from Marsit et al. ^54^ and laboratory isolates were manually detected through the coverage plots of the genomic reads mapping. For all other isolates, the aneuploidy annotations from Peter et al. ^9^ were considered. All strain details including aneuploidy, phylogenetic classification, ecological origin of isolation, and ploidy are provided in SI Table 1.

Mat A disomic yeast strains, constructed by Angelika Amon’s lab ^1^, were kindly provided by Rong Li (Disome WT - strain 11311, Disome 1 - strain 12683, Disome 2 - strain 12685, Disome 4 - strain 24367, Disome 5 - strain 14479, Disome 8 - strain 13628, Disome 9 - strain 13975, Disome 10 - strain 12689, Disome 11 - strain 13771, Disome 12 - strain 12693, Disome 13 - strain 12695, Disome 14 - strain 13979, Disome 15 - strain 12697, Disome 16 - strain 12700).

### Growth curves

To record growth curves, the whole library was revived on agar plates containing synthetic minimal medium (6.8 g/L YNB without amino acids, 2% glucose, 2% agar). Subsequently, colonies were inoculated in synthetic minimal liquid medium (200 μL) and incubated at 30 °C overnight. An aliquot of the yeast culture (2 μL) was transferred to microtiter plates and diluted 100 times in synthetic minimal liquid medium (200 μL total volume per well). Plates were placed into a Spark Stacker (Tecan) in a 30 °C incubation room and the OD at 600 nm was recorded for 48 h. Data analysis was performed in R using the package grofit with default parameters. Strains for which no curve fitting could be performed due to poor growth were omitted from analyses.

### High-throughput yeast culture and lysis for proteomics

#### Natural isolates

The yeast samples were cultivated and digested as follows: the whole collection was grown on agar plates containing synthetic minimal medium (6.8 g/L yeast nitrogen base, 2% glucose, without amino acids). Subsequently, colonies were inoculated in synthetic minimal liquid medium (200 μL) and incubated at 30 °C overnight. 160 µL of the culture were transferred to 96-deep-well plates pre-filled with one borosilicate glass bead in each well and diluted 10 times in synthetic minimal liquid medium to 1.6 mL total volume per well. Plates were sealed using an oxygen-permeable membrane and grown at 30 °C to exponential phase, shaking at 1000 rpm for 8 h. 1.5 mL of cell suspension were transferred to a new deep-well plate and harvested by centrifugation (3,220 x g, 5 min, 4 °C). The supernatant was discarded and plates were immediately cooled on dry ice, then stored at −80 °C until further processing.

#### Disomic strains

Samples were grown using SD-His+G418 agar and medium selecting for the duplicated chromosomes (6.7 g/L yeast nitrogen base without ammonium sulfate, Difco Cat#233520; 20 g/L glucose, Sigma Cat#Y0626; 1 g/L monosodium glutamate, VWR Cat#27872.298; 0.56 g/L CSM-His-Leu-Met-Trp-Ura, MP Biomedicals Cat#4550422; 0.02 mg/mL uracil; 0.06 mg/mL leucine; 0.02 mg/mL methionine; 0.04 mg/mL tryptophan; 200 µg/mL G418, Gibco Cat#11811023). Each disomic strain and the euploid wild type were set up in triplicate. The procedure for cultivation and lysis of the disomic strains was as described above, except for the harvest being conducted at 2700 x g, 10 min, 4 °C.

### Proteomics sample preparation

#### Natural isolates

The samples for proteomics were prepared in 96-well plates as previously described ^32^, with up to four plates processed in parallel. For yeast lysis, 200 µL of lysis buffer (100 mM ammonium bicarbonate, 7 M urea) and ∼100 mg glass beads were added to each well, followed by 5 min bead beating at 1500 rpm (Spex Geno/Grinder). For reduction and alkylation, 20 μL of 55 mM DL-dithiothreitol (1 h incubation at 30 °C) and 20 μL of 120 mM iodoacetamide (incubated for 30 min in the dark at ambient temperature). Subsequently, 1 mL of 100 mM ammonium bicarbonate was added per well, centrifuged (3220 x g, 3 min) and 230 μL of this mixture were transferred to plates pre-filled with 0.9 μg trypsin per well. The samples were incubated for 17 h at 37 °C and the digestion was subsequently stopped by adding 24 μL of 10% formic acid. The mixtures were cleaned-up using C18 96-well plates, with 1 min centrifugations between the steps at the described speeds. The plates were conditioned with methanol (200 μL, centrifuged at 50 x g), washed twice with 50% acetonitrile (ACN, 200 μL, centrifuged at 50 x g) and equilibrated three times with 3% ACN/0.1% formic acid (200 μL, centrifuged at 50, 80 and 100 x g, respectively). Then, 200 μL of digested sample was loaded (centrifuged at 100 x g) and washed three times with 3% ACN/0.1% formic acid (200 μL, centrifuged at 100 x g). After the last washing step, the plates were centrifuged at 180 x g. Subsequently, peptides were eluted in three steps, twice with 120 μL and once with 130 μL of 50% ACN (180 x g), and collected in a plate (1.1 mL, square well, V-bottom). The collected material was completely dried on a vacuum concentrator and redissolved in 40 μL of 3% ACN/0.1% formic acid before transfer to a 96-well plate. The final peptide concentration was estimated by absorption measurements at 280 nm with a Lunatic photometer (Unchained Labs, 2 µL of sample). All pipetting steps were performed with a liquid handling robot (Biomek NXP) and samples were shaken on a thermomixer (Eppendorf ThermomixerC) after each step.

#### Lab-engineered strains

Lysis, reduction, alkylation, and digest of the disomic strains were performed as described above. The digest was quenched using 25 µL of 10% formic acid per sample. The conditioning of the SPE plates was carried out as above, but using 0.1% formic acid instead of the 3% ACN/0.1% formic acid mixture. After loading 200 µL of the digested sample, the columns were washed four times with 200 µL of 0.1% formic acid followed by centrifugation (150 x g). Purified peptides were collected by three consecutive elution steps using 110 µL of 50% ACN (centrifugation at 200 x g). After vacuum drying, peptides were dissolved in 30 µL of 0.1% formic acid. All steps of the sample preparation were performed by hand. Peptide concentrations were determined using a fluorimetric peptide assay kit following the manufacturer’s instructions (Thermo Scientific, Cat#23290).

### LC–MS/MS measurements

#### Natural isolates

For the collection of natural isolates, liquid chromatography was performed on a nanoAcquity UPLC system (Waters) coupled to a Sciex TripleTOF 6600. Peptides (2 μg) were separated on a Waters HSS T3 column (150 mm × 300 μm, 1.8 μm particles) ramping in 19 minutes from 3% B to 40% B (Solvent A: 1% acetonitrile/0.1% formic acid; solvent B: acetonitrile/0.1% formic acid) with a non-linear gradient (SI Table 9). The flow rate was set to 5 μL/min. The SWATH acquisition method consisted of an MS1 scan from m/z 400 to m/z 1250 (50 ms accumulation time) and 40 MS2 scans (35 ms accumulation time) with variable precursor isolation width covering the mass range from m/z 400 to m/z 1250 (SI Table 10).

#### Lab-engineered strains

Proteomics measurements were performed on an Agilent 1290 Infinity LC system coupled to a SCIEX TripleTOF 6600 equipped with an IonDrive source as previously described ^32^. Buffer A consisted of 0.1% formic acid in water, and Buffer B of 0.1% formic acid in acetonitrile. All solvents were LC–MS grade. 5 µg of peptides per sample were separated at 30°C with a 5 min active gradient starting with 1% B and increasing to 36% B on an Agilent Infinitylab Poroshell 120 EC-C18 column (2.1 × 50mm 1.9 μm particle size). The flow rate was set to 0.8 mL/min and the scanning SWATH acquisition method consisted of a m/z 10 wide sliding isolation window..

### Generation of an experimental spectral library for strain S288c

5 µg of yeast digests were injected and run on a nanoAcquity UPLC (Waters) coupled to a SCIEX TripleTOF 6600 with a DuoSpray Turbo V source. Peptides were separated on a Waters HSS T3 column (150 mm × 300 µm, 1.8 µm particles) with a column temperature of 35 °C and a flow rate of 5 µL/min. A 55-min linear gradient ramping from 3% ACN/0.1% formic acid to 40% ACN/0.1% formic acid was applied. The ion source gas 1 (nebulizer gas), ion source gas 2 (heater gas) and curtain gas were set to 15 psi, 20 psi and 25 psi, respectively. The source temperature was set to 75 °C and the ion spray voltage to 5,500 V. In total, 12 injections were run with the following m/z mass ranges: 400–450, 445–500, 495–550, 545–600, 595–650, 645–700, 695–750, 745–800, 795–850, 845–900, 895–1,000 and 995–1,200. The precursor isolation window was set to m/z 1 except for the mass ranges m/z 895–1,000 and m/z 995–1,200, where the precursor windows were set to m/z 2 and m/z 3, respectively. The cycle time was 3 s, consisting of high- and low-energy scans, and data were acquired in ‘high-resolution’ mode. The spectral libraries were generated using library-free analysis with DIA–NN directly from these Scanning SWATH acquisitions. For this DIA–NN analysis, MS2 and MS1 mass accuracies were set to 25 and 20 ppm, respectively, and scan window size was set to 6.

### Ubiquitinomics

Selected aneuploid and euploid yeast isolates were cultivated in SM medium (6.7 g/L yeast nitrogen base with ammonium sulfate, Difco Cat#291920, 20 g/L glucose, Sigma Cat#Y0626) at 30 °C. Three individual pre-cultures per strain were cultured for 16 h, and used to inoculate three flasks of 30–50 mL SM medium per strain. Cultures were harvested at mid-log phase by centrifugation (2880 x g, 8 min, 4 °C) and pellets frozen at –20 °C. Cells were lysed using glass beads (volume equal to pellet volume) in 200 µL of freshly prepared SDC buffer (1% sodium deoxycholate, 10 mM TCEP, 40 mM chloroacetamide, 75 mM Tris-HCl, pH 8.5) by five cycles of 1 min vortexing, 1 min on ice. Samples were centrifuged (13800 x g, 15 min, 4 °C) and the supernatant was collected. Protein concentrations were determined using a Pierce BCA Protein Assay Kit (Thermo Scientific, Cat#23225). 500 µg of proteins were digested with trypsin/Lys-C mix (V5071 or V5072, Promega) overnight at 37 °C with a 1:50 enzyme to protein ratio. K-GG peptide enrichment was performed as reported previously ^40^. The digestion was stopped by adding two volumes of 99% ethylacetate/1% TFA, followed by sonication for 1 min using an ultrasonic probe device (energy output ∼40%). The peptides were desalted using 30 mg Strata-X-C cartridges (8B-S029-TAK,Phenomenex) as follows: a) conditioning with 1 mL isopropanol; b) conditioning with 1 mL of 80% ACN/5% NH_4_OH; c) equilibration with 1 mL of 99% ethylacetate/1% TFA; d) loading of the sample; e) washing with 2 × 1 mL of 99% ethylacetate/1% TFA; f) washing with 1 mL of 0.2% TFA; g) elution with 2 × 1 mL of 80% ACN/5% NH_4_OH. The eluates were snap-frozen in liquid nitrogen and lyophilized overnight. K-GG peptide enrichment was performed by resuspending lyophilized peptides in 1 mL of cold immunoprecipitation (IP) buffer (50 mM MOPS pH 7.2, 10 mM Na_2_HPO_4_, 50 mM NaCl). Peptides were then incubated with 4 µl of K-GG antibody bead-conjugate (Cell Signaling Technology, PTMScan® HS Ubiquitin/SUMO Remnant Motif (K-ε-GG) Kit Cat#59322) for 2 h at 4 °C with end-over-end rotation. Beads were washed (with the help of a magnetic stand) four times with 1 mL of IP buffer and an additional time with cold Milli-Q® water. After removing all the supernatant, the beads were incubated with 200 µL of 0.15 % TFA at room temperature while shaking at 1400 rpm. After briefly spinning, the supernatant was recovered and desalted using in-house prepared, 200 µL two plug StageTips ^55^ with SDB-RPS (3M EMPORE™, Cat#2241). SDB-RPS StageTips were conditioned with 60 µL isopropanol, 60 µL 80% ACN/5% NH_4_OH and 100 µL 0.2% TFA. The K-GG enrichment eluate (0.15% TFA) was directly loaded onto the tips followed by two washing steps of 200 µL 0.2% TFA each. Peptides were eluted with 80% ACN/5% NH_4_OH. Peptides were Speedvac dried and then resuspended in 10 µL of 0.1% FA, of which 4 µL were injected into the mass spectrometer.

For LC–MS measurement, peptides were loaded on 40 cm reversed phase columns (75 µm inner diameter, packed in-house with ReproSil-Pur C18-AQ 1.9 µm resin [ReproSil-Pur®, Dr. Maisch GmbH]). The column temperature was maintained at 60°C using a column oven. An EASY-nLC 1200 system (ThermoFisher) was directly coupled online with the mass spectrometer (Q Exactive HF-X, ThermoFisher) via a nano-electrospray source, and peptides were separated with a binary buffer system of buffer A (0.1% formic acid (FA) plus 5% DMSO) and buffer B (80% acetonitrile plus 0.1% FA plus 5% DMSO), at a flow rate of 300 nL/min. The mass spectrometer was operated in positive polarity mode with a capillary temperature of 275 °C. The DIA method consisted of a MS1 scan (m/z = 300–1,650) with an AGC target of 3 × 10^6^ and a maximum injection time of 60 ms (R = 120,000). DIA scans were acquired at R = 30,000, with an AGC target of 3 × 10^6^, ‘auto’ for injection time and a default charge state of 4. The spectra were recorded in profile mode and the stepped collision energy was 10% at 25%. The number of DIA segments was set to achieve an average of 4–5 data points per peak. For details on the DIA method setup, see Steger et al. ^40^.

### Proteomics data processing

#### Natural isolates

Protein-wise fasta files were created by inferring single nucleotide polymorphisms for each strain based on the reference genome of the S288c strain. In case of heterozygosity, one of the possible alleles was randomly inferred ^9,31^. For non-reference genes, a single representative sequence per protein was available based on the genomes. The proteome for the reference strain S288c was obtained from Uniprot (UP000002311, accessed 10/02/2020). Sequences of strains present in the original strain collections ^9,31^ and subject to intellectual property restrictions were excluded from our study, leading to the inclusion of 1023 strains in the processing.

In order to reduce the processing time and limit the search space to relevant peptides, the protein-wise fasta files were processed to select peptides well shared across the strain collection. The protein sequences were thus trypsin-digested *in-silico* whilst disregarding missed cleavages. Non-proteotypic peptides were excluded and only peptides shared by 80% of the strains were selected for further analysis. This list of peptides was used to filter the experimental library. Raw mass spectrometry files were processed using the filtered spectral library with the DIA-NN software (Version 1.7.12) ^33^. Default parameters of the software were used except for the following: Mass accuracy: 20, Mass accuracy MS1: 12. As the peptides selected were not necessarily present ubiquitously in all the strains, an additional step was required to remove false-positive peptide assignments (entries where a peptide is detected in a strain where it should be absent). This filter led to the exclusion of ∼1% of the entries.

Samples with insufficient MS2 signal quality (∼5.7 × 107), and entries with Q.Value (> 0.01), PG.Q.Value (> 0.01) were removed. Outlier samples were detected based on both the total ion chromatograms (TIC) and the number of identified precursors per sample (Z-Score > 2.5 SD) and were excluded from further analysis. Precursors normalized values as inferred by DIA-NN that were well detected across the samples (in at least 80% of the strains) and with CV < 0.3 in the quality control samples were retained. Subsequently, batch correction was carried out at the precursor level by bringing median precursor quantities of each batch to the same value. Proteins were then quantified using the maxLFQ ^56^ function implemented in the DIA-NN R package, resulting in a dataset containing 1576 proteins for 796 strains. Missing values (<4% of all values) were imputed using K-Nearest Neighbors (KNN) imputation ^57^.

#### Disomes

Mass spectrometry files were processed using the experimental spectral library obtained through gas phase fractionation for the S288c strain with the DIA-NN software (Version 1.7.12). Default parameters of the software were used except for the following: Mass accuracy: 20, Mass accuracy MS1: 12. The output from the software was then processed in R. Entries with Q.Value (> 0.01), PG.Q.Value (> 0.01) and non-proteotypic peptides were removed. Samples with too low OD (< 0.075) were filtered out for further analysis (Disome 4 and one replicate of Disome 8). The precursor normalized values inferred by DIA-NN were used and precursors well detected across 80% of the samples were retained. Proteins were then quantified using the maxLFQ function implemented in the DIA-NN R package. The resulting dataset consists of 1377 proteins for 38 samples. Missing values (< 2.35% of all values) were imputed using the KNN approach. The median value of all available replicate measurements was used for each protein during all further analyses.

#### Ubiquitinomics

Raw data-independent acquisition data files were analyzed using DIA-NN (Version 1.8) in library-free mode searching against the *Sc* reference proteome (strain ATCC 204508/S288c, UniProt ID UP000002311, excluding isoforms, accessed 18.11.2021). Trypsin/P, one missed cleavage, a maximum of two variable modifications (incl. cysteine carbamidomethylation and K-GG) and a precursor charge rate between 2–4 were set for precursor ion generation. The MBR and remove interferences options were enabled and Robust LC (high precision) chosen as the quantification strategy. Peptides carrying a diglycine remnant (UniMod:121) were used to quantify genes using the built-in MaxLFQ algorithm in DIA-NN, with global and run-specific FDR of 1% being applied at both the precursor and the protein group level. The resulting data were filtered to only include genes that were measured in at least two of the three biological replicates in at least one strain for further analyses.

### RNA sequencing

A collection of 1010 unique *Saccharomyces cerevisiae* isolates was gathered from different studies (SI Table 11). 981 isolates came from the 1011 strains collection ^9^, 26 strains from Marsit et al. ^54^, and three lab strains were added: FY4 (a haploid S288C strain), Σ1278b and CEN.PK.

The growth rate of these isolates was estimated by measuring the OD during a 48h-culture in synthetic complete (SC) media with glucose as a carbon source with a microplate reader (Tecan Infinite F200 Pro). Isolates were then grouped according to their growth rates in 96-well plates and grown in SC media until reaching exponential growth phase and an OD of ∼ 0.3. Cells were then transferred in 96-well filter plates, filtered, flashed frozen in liquid N2 and stored at −80°C before mRNA extraction.

A final volume of 10 µL of purified mRNA was obtained using the Dynabeads® mRNA Direct Kit (ThermoFisher, Cat#61012) from an optimized protocol to work in 96-well plates ^42^.

Sequencing libraries were prepared with the NEBNext® Ultra™ II Directional RNA Library Prep Kit for Illumina (NEB, Cat#E7765L) in 96-well plates. 96 combinations of TS HT dual index duplex mixed adapters from IDT® (Integrated DNA Technologies®) were used to barcode the cDNA of each sample from a 96-well plate. The Illumina P5 and P7 universal primers were finally added to the barcoded DNA by PCR. PCR products were then purified and quantified using the Qubit™ dsDNA HS Assay Kit (Invitrogen™) in a 96-well plate with a microplate reader (Tecan Infinite F200 Pro) with the excitation laser set at 485 nm and the emission laser at 528 nm. 20 ng of DNA from each sample of a 96-well plate were pooled to be sequenced. After quantity and quality controls, each pool was sequenced on Nextseq 550 high-output at the EMBL Genomics Core Facilities. A mean of 6.45 million single-end reads of 75 bp was obtained for each sample after demultiplexing.

### RNAseq processing

Raw reads were cleaned with cutadapt ^58^ to remove adapter as well as low quality reads that were trimmed on the basis of a Phred score threshold of 30 and discarded if less than 40 nt long after this trimming step. For each of the isolates, clean reads were mapped to the *Sc* reference sequence in which the SNPs of the corresponding strains were inferred (as described in ^9^) plus the accessory genes that were not classified as ancestral or *S. paradoxus* orthologs in Peter et al. (^9^, n = 395). After the mapping step using STAR ^59^, isolates with more than 1 million reads mapped were kept for analysis, resulting in a final set of 969 strains (SI Table 11). Mapped reads counts were then obtained using the featureCounts function from the Subread package ^60^ with the genes described in the *Sc* reference annotation (n = 6,285) and accessory genes (n = 395) as features.

Raw read counts were filtered to only include genes with a mean of more than 1 count per million (CPM) across measured strains. These filtered read counts were then normalized using the trimmed-mean of M-values method ^61^ as implemented in edgeR ^62,63^. Non-zero, non-log_2_-transformed counts-per-million values were used for further analysis.

### Bioinformatic analysis

All bioinformatic analyses were conducted in R 3.6 unless otherwise indicated. KEGG annotations for *Sc* genes were obtained through the KEGG database (accessed 15.01.2021). The org.Sc.sgd.db package ^64^ was used to obtain chromosomal location information for genes and to map gene names to systematic ORF identifiers. If no gene name was annotated in this package, the systematic ORF identifier was used instead. Standardized *Sc* yeast strain names and systematic ORF identifiers for genes were used throughout all analyses.

#### Assembly of integrated chromosome copy number, mRNA, and protein expression dataset

Chromosome copy number status for all engineered disomic strains was confirmed by Torres et al. ^1^, meaning all disomic strains used in our study were haploid with indicated “disomic” chromosomes duplicated. One exception was Disome 13: despite published mRNA expression values being available and proteomics data having been measured in our experiments, Disome 13 was excluded from all analyses due to having undergone whole-genome duplication when reaching our laboratory (private communication, J. Zhu). Gene copy numbers for natural strains were downloaded from the 1002 Yeast Genome website (http://1002genomes.u-strasbg.fr/files/) ^9^. Only genes identified with a systematic ORF ID and with non-zero and non-missing values were used in further analyses. Microarray gene expression data for disomic strains were downloaded from the supplementary material of the work describing the disomic strain collection ^1^. Only data for strains grown in batch culture were used. Raw expression profiling data for this dataset are available from the GEO database under the accession number GSE7812.

Chromosome copy numbers of the induced meiosis stable aneuploid strains were used as described by Pavelka et al. ^2^, and proteomics data were downloaded from the SI material of the same publication.

Data for gene copy number, mRNA expression, and protein abundances were matched by strain name and systematic ORF identifier for both the disomic strain collection and the natural isolate collection. Only genes for which values for gene copy number, transcript, and protein levels were available were used for analyses. We noticed that a number of strains in the natural isolate collection exhibited a mismatch between the median gene copy number per chromosome and the assigned aneuploidy as described in SI Table 1, most likely attributable to segmental aneuploidies and algorithm-specific thresholds used for aneuploidy determination. We excluded all strains containing one or more of those “mismatched” chromosomes from our analysis (SI Table 3). From this point, chromosome copy numbers as given by the aneuploidy annotation were used throughout the analyses.

#### Comparative mRNA and protein expression analysis

The assembled integrated dataset was used to compare the relative changes in chromosome copy number, mRNA transcript expression, and protein abundances between aneuploid and euploid strains.

Relative chromosome copy number changes were calculated as the log_2_ ratio between chromosome copy number and ploidy of the strain. Relative abundances for transcriptomic and proteomic data were calculated gene-wise as the log_2_ ratio between a gene’s mRNA or protein abundance in a given strain and the median mRNA or protein expression value of the respective gene across all euploid strains. For the transcriptomic data of the lab-engineered disomic strains, we used the transcript levels as published, meaning as log_2_ fold-changes compared to the wild-type strain (Disome WT, 11311) ^1^. In case of replicate measurements, the median value was used for further analysis. For the proteomic data of the aneuploids generated by induced meiosis of triploid and pentaploid strains, we used the published replicate-averaged normalized spectral abundance factors (dNSAF) ^2^, including only proteins that were quantified in at least 2 out of 4 replicates in at least one strain, and calculated log_2_ protein ratios as described above with strain RLY2626 being the euploid reference.

For the log_2_ mRNA and protein ratios of genes encoded on euploid chromosomes, a distribution centered around 0 would be expected, representing no overall shift of relative expression values of these genes across strains. This was true for our proteomics data, and also for most strains in the transcriptomic data. Since some natural isolates showed left tails in this distribution for the transcriptomic data, presumably because of restricting the assembled dataset to genes for which we had data across all three -omics layers, we decided to normalize the relative mRNA and protein expression values. Normalization was carried out in a strain-by-strain manner. First, the median log_2_ mRNA or protein ratio of all genes encoded on euploid chromosomes of a given strain was calculated. This median value was then subtracted from all log_2_ mRNA or protein ratios of that strain. To assess buffering at the chromosome level, the median mRNA or protein log_2_ ratio of all genes encoded per chromosome was calculated.

The proteome profiles of Disome 12 and Disome 14 showed no aneuploid signature, indicating that those strains, even though they were held under selective pressure, had lost their duplicated chromosome either before their arrival in our laboratory or during our experiments (Fig S5). Both strains were therefore excluded from our analysis. Similarly, when comparing relative chromosome copy number changes and relative mRNA expression levels in the natural isolate collection, we noticed discrepancies indicative of chromosomal instabilities in natural *Sc* isolates. Some euploid strains had gained or lost chromosomes, evident as much higher or lower fold-change of expression values observed in the transcriptomics data than in chromosome copy numbers. Likewise, some aneuploid strains underwent changes in their karyotype, resulting in either more complex aneuploidies, or in aneuploid strains reverting to euploid strains. We decided to include in our analysis only strains that showed consistent relative expression (log_2_ ratio) values on the chromosome copy number and the transcriptome level. Consequently, we excluded strains that had at least one chromosome for which the difference between relative chromosome copy number and the median of the normalized relative mRNA abundances differed by more than ±4 standard deviations from the mean, based on all relative chromosome–mRNA comparisons (SI Table 4). After exclusion of these strains, the calculations to obtain gene-wise relative (strain/euploid) mRNA and protein expression values were repeated to avoid unintended biases towards these excluded strains.

#### Quantification of chromosomal buffering of aneuploidy

For both disomic lab-engineered strains and natural isolates of *Sc*, mRNA and protein expression log_2_ ratios between a gene’s expression in a strain and the gene’s expression over all euploid strains were examined in relation to the log_2_ fold change of the copy number of the chromosome the gene is located on. For gains of chromosomes, all log_2_ chromosome copy number changes for which less than 300 affected data points (genes) were quantified were excluded for distribution visualization. For chromosome losses, fewer data points were available overall, so the described cut-off was set at 50 data points (genes). To quantify the relationship between chromosome gains (log_2_ chromosome copy number/ploidy > 0, all relative chromosome copy number changes included) and relative mRNA or protein expression from aneuploid chromosomes, linear models were fitted between the log_2_ chromosome copy number change and the median relative mRNA or protein expression value.

#### Assessment of the global proteome response in aneuploid strains

To find genes differentially expressed in *trans* in aneuploid strains, i.e. genes encoded on euploid chromosomes of aneuploid strains that show up-or downregulation at the mRNA or protein level when compared to euploid strains, we calculated the gene-by-gene median normalized relative mRNA or protein abundances (log_2_ ratios) of all genes encoded on euploid chromosomes in aneuploid strains. KEGG pathway gene set enrichment analysis of these median relative expression values was conducted using Webgestalt 2019 using default settings (accessed online 01/12/2021) ^65^. Additionally, non-parametric one-sample Wilcoxon tests were used to compare gene-by-gene median normalized protein log_2_ ratios across euploid chromosomes of aneuploid strains against the theoretical gene-by-gene median protein log_2_ ratio value across euploid strains. P-values were corrected for multiple hypothesis testing using the Benjamini-Hochberg method as implemented in the rstatix package ^66^.

#### Determination of ubiquitination levels

Relative levels of ubiquitinated proteins were determined gene-wise by calculating the log_2_ ratio between the measured abundance of a ubiquitinated protein in each strain and the median abundance of the ubiquitinated protein across all euploid strains. Assuming a distribution centered around 0 of relative levels of ubiquitinated proteins on euploid chromosomes, these relative abundances were then normalized strain-wise by subtracting the calculated median log_2_ ratio of all genes expressed on euploid chromosomes of a strain from all log_2_ ratios of that strain.

### Proteasome activity assays

Biological replicates of aneuploid and euploid yeast strains were cultivated, harvested, and frozen as described for the ubiquitinomics experiments. Using a volume of glass beads (425–600 µm, Sigma #G8772) equal to the size of the cell pellet, cells were resuspended in 200 µL non-denaturing lysis buffer (20 mM HEPES-KOH pH 7.4, 110 mM potassium acetate, 2 mM magnesium chloride, 1 mM EDTA pH 7.0, 2 mM ATP) and lysed by five cycles of 1 min vortexing, 1 min on ice. Samples were subsequently centrifuged at 21100 x g for 15 min. 150 µL of the supernatant were aspirated well above the pellet and transferred to a new tube in order to avoid interferences from beads and cellular debris in the assay. The protein concentration of the samples was measured using a Pierce BCA Protein Assay Kit (Thermo Scientific, Cat#23225). To determine the proteasome activity, a fluorogenic substrate was used (Suc-Leu-Leu-Val-Tyr-AMC, Bachem ## I-1395, prepared as a 20 mM stock solution in DMSO). Turnover of the substrate is a proxy for the chymotryptic activity of proteasomes and results in an increase of fluorescence emission. Samples were diluted to 0.5 µg/µL in assay buffer (50 mM Tris pH 7.5, 5 mM magnesium chloride, 1 mM ATP), and the substrate was diluted to 1.11 µM/µL in assay buffer. 20 µL of diluted sample and 50 µL of substrate were mixed in black 96-well microtiter plates and incubated for 30 min at 37 °C. Samples were measured using a Bio-Tek Synergy HT with 360 nm excitation, 460 nm emission. For each sample, three technical replicates were measured and averaged. All measurements were background-corrected. Proteasomal activity for genetically close aneuploid and euploid strains was always measured in the same assay batch. The activity of euploid samples was calculated relative to the mean intensity across samples of the genetically close aneuploid strain.

## Data availability

Raw and processed mass spectrometry data will be released upon publication.

## Code availability

Custom analysis scripts are available upon request.

## SI Figures

**SI Fig. 1:**
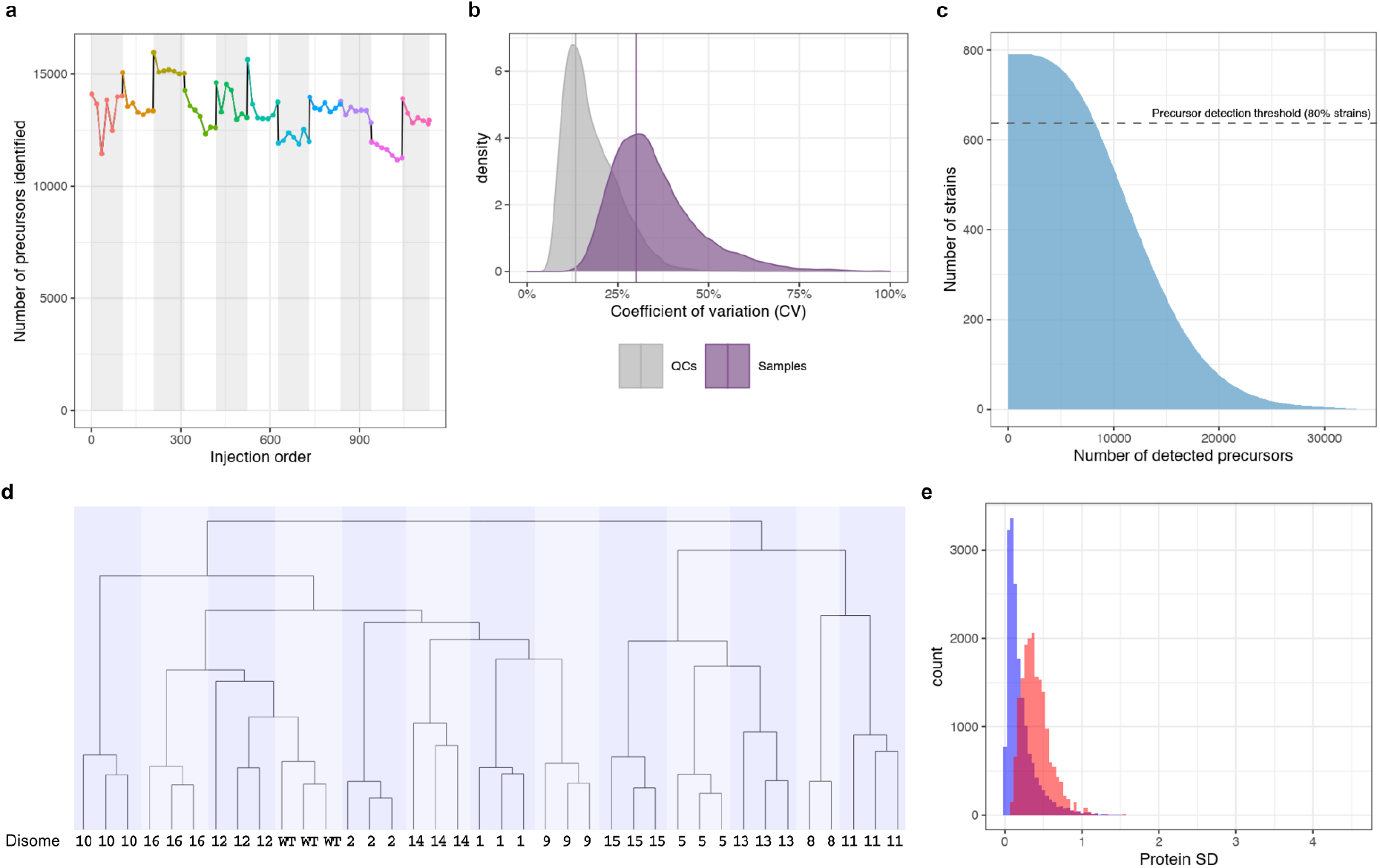
High-throughput proteomics platform and quality assessment plots for proteomics data processing. (a) Number of precursors identified by DIA-NN in the samples of the natural isolate collection, before further processing, ordered by injection number, demonstrating a stable performance over the acquisition of the data. Colors and shaded backgrounds highlight separate batches. (b) Coefficients of variation (CV) of the precursor quantities in the technical quality controls (QCs, technical variability, n= 77) compared to the biological samples (yeast isolates, biological signal, n= 796). The solid purple line indicates the short, a robust estimator of the interval covering half of all values ^67^ (c) Precursors detection rate in the natural isolates. A strict threshold was set to retain precursors that were well detected in at least 80% of the strains. The precursors retained were then used for protein quantification. (d) Dendrogram of the hierarchical clustering of the disome replicates based on their proteomes. Numbers indicate the name of the disomic strain, WT is the haploid euploid parental strain (W303). (e) Variance within replicates vs. variance between replicates of the disomic strain proteomics measurements demonstrating low technical variability and a well-detected biological signal in the disome dataset. The comparison of the standard deviations between replicates (blue) and when considering all samples (red) is shown.

**SI Fig. 2:**
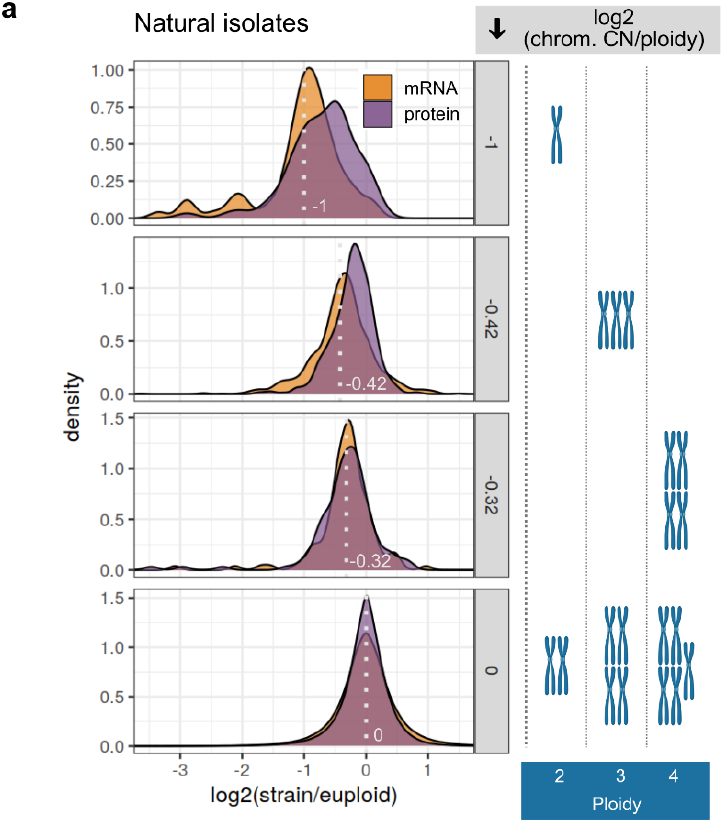
(a) Distributions of log_2_ protein (purple) and mRNA (orange) ratios across all euploid or aneuploid chromosomes of aneuploid natural strains that show chromosome losses. Genes are binned according to the relative copy number change of the chromosome they are encoded on. The ploidy of the strains included in the analysis is indicated (blue box; 2: diploid, 3: triploid, 4: tetraploid), and absolute chromosome numbers leading to the shown relative chromosome copy number changes are depicted to the right of each panel. The light gray dotted lines and numbers denote the relative chromosome copy number change (log_2_ chrom. CN/ploidy) in the density plots. For all distributions, relative expression levels are shown between −3.5 and 1.5.

**SI Fig. 3:**
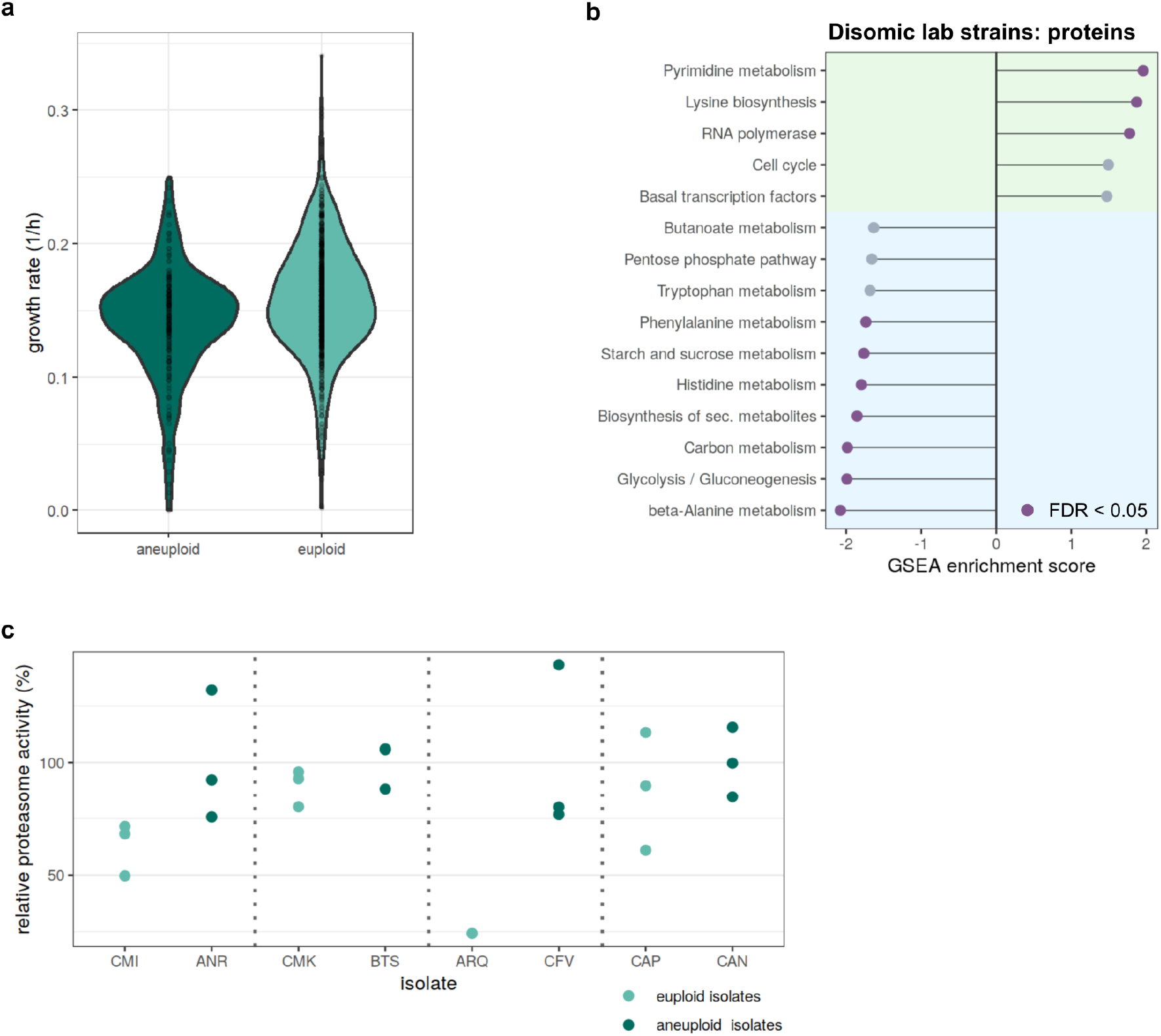
(a) Growth rates of aneuploid and euploid isolates for which proteomics data were collected. Strains were cultivated in synthetic minimal medium. See also SI Table 2 for all growth parameters. (b) Gene set enrichment analysis of median log_2_ protein expression ratios (strain/euploid) of genes encoded on euploid chromosomes across all disomic strains. Statistically significant enrichment scores (false discovery rate <0.05) are colored in purple (protein). (c) Proteasome activity and ubiquitination in aneuploid vs. euploid natural *Sc* isolates. Chymotryptic activity of proteasomes was measured in native cell lysates of pairs of genetically close natural strains. Strains are identified by their standardized isolate name (SI Table 1). Experiments were performed as biological triplicates (except for ARQ) with three technical repeat measurements per sample. The proteasome activity of each sample is shown relative to the mean intensity of all measurements of the respective genetically close aneuploid strain.

**SI Fig. 4:**
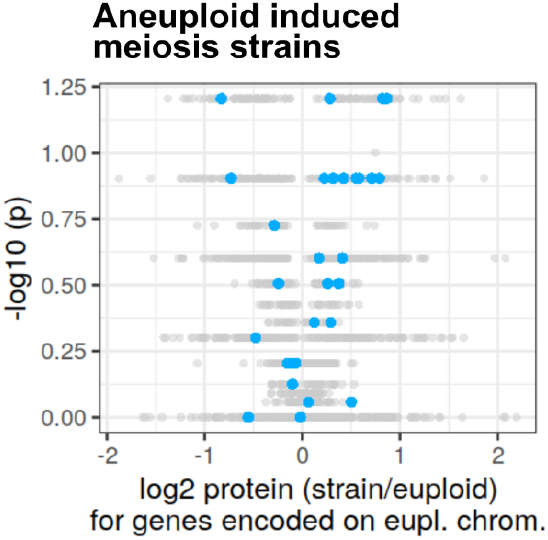
Volcano plot for induced meiosis strains showing results of non-parametric one-sample Wilcoxon tests comparing the median log_2_ protein ratios to the theoretical gene-by-gene median protein log_2_ ratio across all euploid strains of the respective collection (µ = 0). Proteins with statistically significant differential expression after multiple hypothesis correction (Benjamini-Hochberg) are colored in dark gray. Structural components of the proteasome are highlighted in blue.

**SI Fig. 5:**
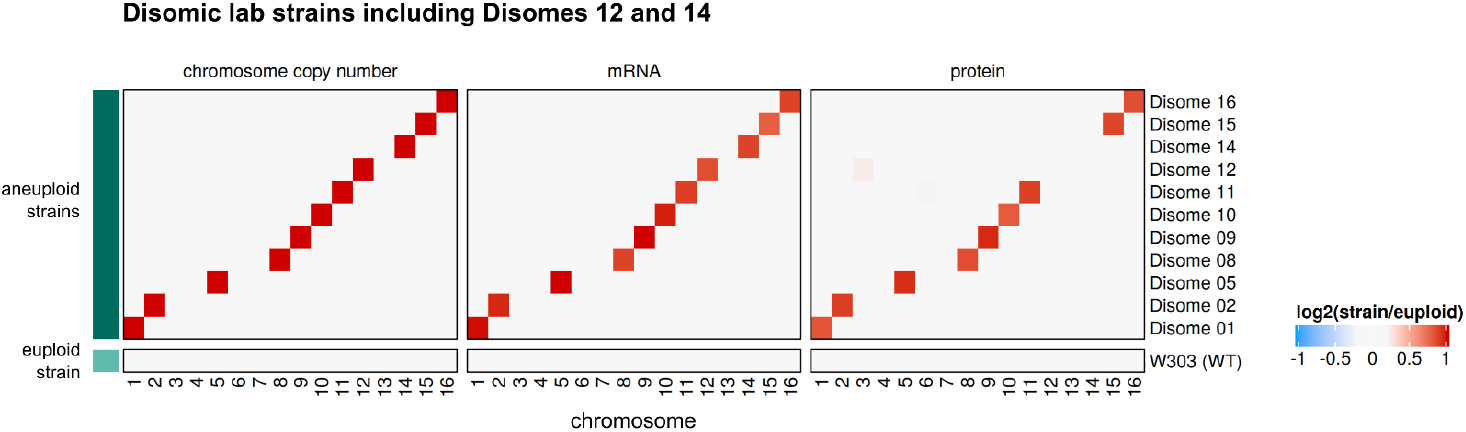
Relative chromosome copy numbers, relative median mRNA, and relative median protein abundances per chromosome between strains and euploid reference (log_2_ ratios strain/euploid) for laboratory-engineered disomic strains including Disomes 12 and 14 that had lost their aneuploid chromosome. Strain W303 (haploid euploid) was used as the reference for calculation of log_2_ ratios.

## Acknowledgements

We would like to thank Pedro Beltrao, Gilles Fischer, and Vadim Farzdinov for helpful discussions, and Hezi Tenenboim for critically reading the manuscript. Jin Zhu, Sreekumar Ramachandran and Rong Li kindly sent us the disomic strains. We thank the Core Facility High Throughput Mass Spectrometry of the Charité for support in sample preparation, acquisition and analysis. Elements of figures were created with BioRender.

This project has received funding from the European Research Council (ERC) under the European Union’s Horizon 2020 research and innovation programme (grant agreement No 951475 - ERC Synergy Award FungalTolerance to J.B. and M.R., PhenomeNal ERC Consolidator grant 772505 to J.S.), from the Ministry of Education and Research (BMBF) as part of the National Research Node ‘Mass spectrometry in Systems Medicine (MSCoreSys) under grant agreement 031L0220 (to M.R.), the Wellcome Trust (Investigator Award IA

200829/Z/16/Z to M.R.), from the BMBF grant 161L0221 (to V.D.), from Fondation pour la Recherche Médicale (EQU202003010413) and Agence Nationale de la Recherche (ANR-15-IDEX-01) (to G.L). This work was further supported by the Francis Crick Institute, which receives its core funding from Cancer Research UK (FC001134), the UK Medical Research Council (FC001134), and the Wellcome Trust (FC001134).

## Author contributions

J. M. analyzed and interpreted data, designed and conducted proteomics experiments with disomic strains, performed ubiquitinomics and proteasome activity experiments.

P. T. developed the raw mass spectrometry data processing pipelines, analyzed the data, and provided scientific input.

F. Ag. conducted proteomics experiments with the natural strain collection, recorded and analyzed growth curves, and provided scientific input.

C. M. designed the high-throughput yeast proteomics sample cultivation and preparation pipeline, performed mass spectrometry measurements for the natural strain collection, and provided scientific input.

A. L. performed sample lysis for ubiquitinomics experiments and performed proteasome activity assays.

M. S. prepared the ubiquitinomics samples for mass spectrometry, conducted ubiquitinomics measurements, and provided scientific input.

T. G. performed genomic analyses, interpreted data and provided scientific input.

E. C. performed RNA sequencing experiments of the strain collection and processed the RNAseq data.

A.-S. E. and F. Am. performed mass spectrometry measurements for the natural strain collection and the disomic strain collection, respectively.

N. B. performed experiments.

V. D.contributed to the data processing pipeline and provided scientific input.

M. M. supervised mass spectrometry measurements for the disomic strain collection and provided scientific input.

M. d. C., G. L., J. S. and J. B. provided scientific input.

M. R. acquired funding, supervised the project, interpreted data. J.M., J.B. and M.R. wrote the manuscript with input from all authors.

